# Development of a diamide resistance diagnostic method using LAMP based on a resistance-specific indel in ryanodine receptors for *Spodoptera exigua* (Lepidoptera: Noctuidae)

**DOI:** 10.1101/2020.05.19.103507

**Authors:** Juil Kim, Hwa Yeun Nam, Min Kwon, Ji Hye Choi, Sun Ran Cho, Gil-Hah Kim

## Abstract

Recently, resistance to diamide insecticides (IRAC group 28) has been reported in various lepidopteran pests, including *Spodoptera exigua*. In the present study, susceptibility of six field populations was evaluated to two diamide insecticides: chlorantraniliprole and flubendiamide. The bioassay test for resistance revealed a high level of diamide resistance and helped to select a diamide resistant (Di-R) strain, whose LC50 values against chlorantraniliprole and flubendiamide were 28,950- and 135,286-fold higher, respectively, than those of susceptible strains. In the ryanodine receptor, instead of the G4946E mutation, one of the well-known diamide resistance mechanisms, we found a I4790M mutation and identified the resistance allele-specific indel linked to it. Resistance allele diagnostic primers were designed using this distinct region and applied in loop-mediated isothermal amplification (LAMP) and general PCR. LAMP accurately detected the specific indel when conducted for 2 h at temperature range from 63 °C to 65 °C and using four LAMP primers; its efficiency was further amplified by an additional loop primer. A broad range of DNA concentrations was workable in the LAMP assay, with the minimum detectable DNA concentration of 100 pg. The new DNA releasing method used for the LAMP assay consisted of 5 min of incubation of a larva or adult tissue at 95°C. The entire diagnostic process, which included the DNA releasing technique and LAMP, lasted only 100 min. This simple and accurate LAMP assay can be applied to monitor diamide resistance and for integrated resistance management of *S. exigua* in the field.

## Introduction

Diamide insecticides are one of the major class of insecticides targeting ryanodine receptors (RyRs) that have been introduced to the market to control a broad range of herbivorous pests, particularly those in the order Lepidoptera (Nauen 2006; Sattelle 2008). The anthranilic diamides, including chlorantraniliprole and cyantraniliprole, were discovered and developed commercially by Dupont (Cordova et al. 2005; Lahm et al. 2005; Froster et al. 2012; Liu et al. 2018). A third systemic anthranilic diamide, cyclaniliprole, was developed by Ishihara Sangyo Kaisha (ISK) biosciences corporation in 2004. The phthalic acid diamide flubendiamide was discovered by Nihon Nohyaku and co-developed with Bayer (Tohnishi et al. 2005). These insecticides were commercialized and rapidly gained market share exceeding US $1.4 billion, accounting for approximately 8% of the insecticide market in 2013 (Sparks and Nauen 2015).

The target site of diamide RyR is the largest known ligand-gated calcium channel found in the sarcoplasmic/endoplasmic reticulum membrane in muscle and nervous tissue (Sun & Xu, 2019). RyR controls the release of calcium from intracellular stores and regulates a variety of cellular and physiological activities, such as gene expression, neurotransmitter release, hormone secretion, muscle contraction, cell proliferation, and finally insect death (Coronado et al. 1994; Nauen and Steinbach 2016). Diamides are broad spectrum insecticides, affecting lepidopterans, with a high safety profile (Jeanguenat, 2013). Because of these advantages, diamides are widely used worldwide. However, after only a few years of application, field-evolved resistance to diamides, such as chlorantraniliprole, has been reported in several lepidopteran pests including *Spodoptera exigua* (Lai et al. 2011), *S. frugiperda* (Boaventura et al. 2020), *Plutella xylostella* (Troczka et al. 2012), and *Tuta absoluta* (Roditakis et al. 2015). The target-site mutation conferring the amino acid substitution G4946 was first identified in a diamide-resistant strain of diamondback moth *P. xylostella* from the Philippines and Thailand (Troczka et al. 2012), and subsequently detected in field populations in many other countries (Guo et al. 2014a, 2014b; Steinbach et al. 2015). Three additional substitutions (I4790M, E1338D, and Q4594L) associated with diamide resistance were found in a field population of *P. xylostella* collected from Yunnan Province of China (Guo et al. 2014a). Diamide resistance mechanism was functionally confirmed by recombinant expression of mutant RyR variants stably expressed in Sf9 cells (Troczka et al. 2015), and by CRISPR/Cas9 genome editing in transgenic *Drosophila melanogaster* and *S. exigua* carrying RyR I4790M and G4946E mutations, respectively (Douris et al. 2017; Zuo et al. 2017). Although the metabolic resistance mechanisms to diamides in *S. exigua* remain largely unknown (Nauen and Steinbach 2016), there is little doubt that it is conferred by a mutation in RyR.

The beet armyworm, *Spodoptera exigua* (Hübner), is a pest of global importance in cultivation of numerous crops, including cotton, tomato, lettuce, cabbage, and ornamentals (CABI 2020). Frequent applications of diamide in Korea selected for field strains of *S. exigua* with high levels of diamide insecticide resistance, exceeding 10,000-fold when compared with that of a susceptible reference strain. However, the G4946E mutation was not detected in all tested resistant strains and/or populations (Cho et al. 2018). Therefore, we examined the toxicity of two major diamide insecticides (chlorantraniliprole and flubendiamide) to six field populations of *S. exigua* larvae to identify populations susceptible to the insecticides. We aimed to identify additional mutations and a resistance-specific indel in RyR in diamide-resistant strains and populations. We also developed a molecular diagnostic method to detect diamide resistance in the field that can be used for more effective pest resistance management and sustainable control of *S. exigua*.

## Materials and methods

### Insects

Eleven field populations of *Spodoptera exigua* were collected in significantly damaged potato, cabbage, and green onion fields from May to August in 2019. These comprised three sites in Gangwon-do (Hongcheon, Gangneung, Hoengseong), three sites in Gyunggi-do (Yeoju, Anseong, Icheon), one site in Chungcheongbuk-do (Cheongju), one site in Gyeongsangnam-do (Miryang), and two sites in Jeollanam-do (Haenam, Jindo) (Fig. 1a). We collected larvae from heavily damaged plants, and some adults were captured using pheromone traps. Six populations were used for bioassay test of resistance and genotyping: Anseong (AS), Cheongju (CJ), Gangneung (GN), Icheon (IC), Jindo (JD), and Yeoju (YJ). Five additional populations were used for genotyping: Haenam (HN), Hoengseong (HS), Hongcheon (HC), Miryang (MY), and Pyeongchang (PC). For bioassays, we collected more than 1,000 larvae of *S. exigua* to calculate the insecticide resistance ratio based on the first generation of indoor breeding. The larvae were collected from two different sites in each region (Jindo, Pyeongchang, Hoengseong, Gangneung, and Hongcheon), and collections from the two sites were pooled into a single population. Two susceptible strains were used. One was reared at Chungbuk National University without insecticide exposure (Cho et al. 2018; thereafter referred to as CNU strain) and used in most experiments; the other susceptible lab strain was obtained from the National Institute of Agricultural Sciences (thereafter, NAS strain) and used as a reference strain in genotyping and molecular diagnosis. A diamide-resistant strain (Di-R) was selected among individuals of Jindo populations that survived two treatments with a recommended concentration of chlorantraniliprole (25 ppm) or flubendiamide (100 ppm). This selection was repeated for four generations. Di-R and the susceptible CNU strain were used as basic strains for elucidating resistance mechanisms. A F1 hybrid was produced by single pair mating of a resistant and susceptible strain (15 pairs of R♂ × S♀ and S♂ × R♀ each). Selected diamide-resistant strains of *S. exigua* and field-collected resistant populations were raised in both the Highland Agriculture Research Institute and Chungbuk National University on an artificial diet (Cho et al. 2018).

**Fig. 1.**
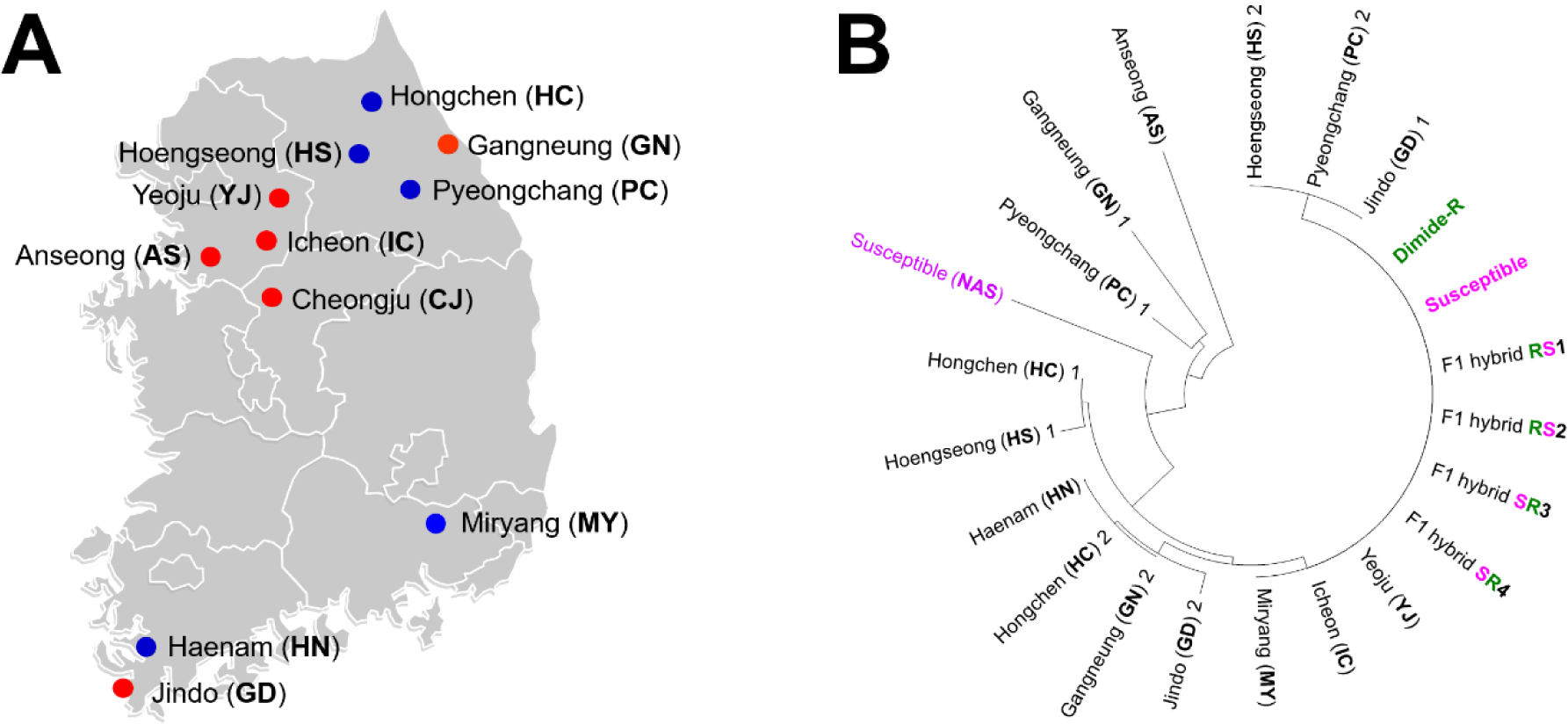
Local populations of *Spodoptera exigua* (A) collection sites and their (B) phylogenetic relationship with two susceptible strains (Susceptible and Susceptible NAS) and diamide resistant strain (Diamide-R) based on partial sequences of the mitochondrial genome. In total 11 local populations collected in 2019 were used. Six populations, indicated by red dots, were used for bioassay test of resistance and genotyping: Anseong (AS), Cheongju (CJ), Gangneung (GN), Icheon (IC), Jindo (JD), and Yeoju (YJ). Five additional populations, indicated with blue dots, were used for genotyping: Haenam (HN), Hoengseong (HS), Hongchen (HC), Miryang (MY), and Pyeongchang (PC). The phylogenetic relationship was inferred from 22 single nucleotide polymorphisms of the 2,544 nucleotide sequences by using the maximum likelihood method based on the Tamura-Nei model.

### Chemicals

Two types of diamide insecticides were used, commercial insecticides (chlorantraniliprole WG and flubendiamide SC) and technical grade insecticide (>99% pure; Sigma-Aldrich, Saint Louis, MO, USA). The former was used in the selection of resistant strain, and latter for bioassay test of resistance.

### Bioassay test of resistance

For the bioassay, six populations collected in the field (AS, CJ, GN, IC, JD, YJ) were reared on artificial diet for at least one generation post-procedure in the lab to obtain third instar larvae. Technical-grade insecticides were dissolved in acetone to 10,000 ppm concentration and dissolved in distilled water mixed with triton (0.2%) to appropriate concentrations. A three-piece artificial diet (a 3 cm^3^ cube) was immersed in insecticide for 30 s and transferred into a Petri dish (10 cm diameter, 4 cm height). Afterward, 10 *S. exigua* larvae were transferred into the Petri dish. All experiments were performed with four replicates. Mortality rate was determined after 72 h of treatment, and the value was used to calculate median lethal concentration (LC50) using the SAS program based on probit model (SAS Institute 9.1, Cary, NC, USA). The resistance ratio was calculated by dividing the LC50 value of the population by that of the CNU susceptible strain.

### Phylogenetic relationship

About 2.5 kb partial sequences of mitochondrial DNA were analyzed to identify the genetic characteristics of each strain. Total DNA was extracted from three individuals randomly selected from each collection site to represent the region by using DNAzol, and then amplified with KOD FX polymerase (Toyobo Life Science, Osaka, Japan) and BAW_mtC2_F and BAW_mtC2_nR3 primer sets. PCR was conducted under the following thermal conditions: initial denaturation at 95 °C for 2 min, followed by 35 cycles of 95 °C for 20 s, 57 °C for 20 s, and 68 °C for 90 s, and final extension for 60 s at 68 °C. The PCR products were directly sequenced (Macrogen, Seoul, Korea). The PCR product was about 2.5 kb, and the entire sequence was analyzed by primer walking using LCO1490 and HCO2198 primers. After contig assembly, sequences were analyzed by Gene Runner ver. 6.5.52 (www.generunner.net), and molecular phylogenetic analysis of the mitochondrial genome was inferred by using a maximum likelihood method implemented in MEGA 7 with bootstrapping (Sanderson and Wojciechowski 2000; Kumar et al. 2016).

### Mutation survey in ryanodine receptors

Total RNA was isolated from five larvae (3rd instar) of susceptible and Di-R strains using a RNeasy Mini Kit according to the manufacturer’s protocol (Qiagen, Hilden, Germany). cDNA was synthesized using a TOP script™ cDNA synthesis kit (Enzynomics, Daejeon, Korea). Genomic DNA (gDNA) was extracted by DNAzol (Molecular Research Center, Cincinnati, OH, USA) from each strain and five larvae (3rd instar) of the 11 populations. cDNA and gDNA samples were quantified by using a Nanodrop (NanoDrop Technologies, Wilmington, DE, USA).

Partial fragments of RyR were amplified using the appropriate primer sets (Table 1) with KOD FX polymerase (TOYOBO). Primers RyR_4790UF and RyR_mU_R were used to verify G496E and I4790M mutations, and primers RyR_4790UF and RyR_I4790M_R or RyR_PASA_R3 were used to identify only the I4790M mutation. PCR was conducted under the following thermal conditions: initial denaturation at 95 °C for 2 min, followed by 35 cycles of 95 °C for 20 s, 57 °C for 20 s, and 68 °C for 90 s, and a final extension at 68 °C for 60 s. The PCR products were directly sequenced (Macrogen). Contig assembly and amino acid sequence alignment were conducted using Lasergene v14 (DNASTAR, Madison, WI, USA) with the neighbor-joining method (Saitou and Nei 1987).

**Table 1.** Susceptibility of susceptible and diamide-resistant (Diamide-R) strains and six local populations of *Spodoptera exigua* to two diamide insecticides

| Strains | Chlorantraniliprole |  | Flubendiamide |  |
| --- | --- | --- | --- | --- |
|  | LC<sub>50</sub>(mgL<sup>-1</sup>)<br>(95% CL)<sup>a</sup> | RR<sup>b</sup> | LC<sub>50</sub>(mgL<sup>-1</sup>)<br>(95% CL)<sup>a</sup> | RR<sup>b</sup> |
| Susceptible | 0.002 | 1 | 0.0007 | 1 |
| Diamide-R | 57.9 (35.4 - 89.7) | 28,950 | 94.7 (79.2–118.9) | 135,286 |
| Anseong | 8 (5.3–12.5) | 4,000 | 0.3 (0.2–0.5) | 428 |
| Cheongju | 1.2 (0.3–2.7) | 600 | 10.5 (7.0–14.4) | 14,957 |
| Gangneung | 6.6 (5.3–8.2) | 3,300 | 210.1 (71.7–295.1) | 300,143 |
| Icheon | 4.6 (2.3–7.0) | 2,300 | 52.31 (32.1–70.0) | 74,729 |
| Jindo | 13.4 (7.6–25.3) | 6,700 | 27.9 (24.1–32.2) | 39,929 |
| Yeoju | 21.2 (9.9–498.0) | 12,500 | 90.4 (67.8–132.0) | 129,186 |

<sup>a</sup> CL: Confidence limits.
<sup>b</sup> Resistance ratio = LC<sub>50</sub> of the resistant strain or local populations/LC<sub>50</sub> of the susceptible strain.

### Diagnostic primer design for lamp loop-mediated isothermal amplification (LAMP) and multiplex PCR

The analysis of aligned partial sequences of susceptible and resistant strains confirmed the existence of a resistance-specific indel (GenBank accession number of susceptible strain, MT292621 and that of Di-R, MT292620). Based on the partial genomic RyR sequence alignment results, the partial sequences containing the resistant-specific indel were re-aligned to design LAMP primers in PrimerExplorer V5 (Eiken Chemical Co., Ltd., Tokyo, Japan). Forward and backward internal primers (FIP and BIP) with backward 3 (BAW_RyR_B3) were designed using the program with some modifications. The main diagnostic primer forward 3 (BAW_RyR_F3) was manually designed inside the resistance-specific region. The forward internal primer (FIP) was a combination of F1c and F2, and the backward internal primer (BIP) was a combination of B1c and B2. Loop primers (BAW_RyR_LF and BAW_RyR_LB) were located between F1c and F2 or between B1c and B2. Multiplex PCR primers were designed manually using partial RyR sequences. A partial *ace1* type acetylcholinesterase gene sequence (DQ280488) was used to design the primers BAW_ace1_D1F and BAW_ace1_D1R, which were used for positive control in multiplex PCR.

### LAMP and multiplex PCR protocol

The general protocol of LAMP was conducted in a 25 μL reaction mixture using a WarmStart® LAMP Kit (New England Biolabs, Ipswich, MA, USA) following the manufacture’s guideline (Kim et al. 2020). To test the DNA releasing technique for insect tissue, approximately 10 mg of larval tissue or an adult leg or antenna was incubated at 95 °C for 5 min with 30 μL nuclease-free water, and 2 μL of the resulting supernatant was then used in the LAMP assay with each primer set.

For multiplex PCR, susceptible and resistance alleles in RyR, as well as partial *ace1*, were amplified using the appropriate primer sets (Table 1) with KOD FX polymerase (TOYOBO). PCR was conducted under the following thermal conditions: initial denaturation at 95 °C for 2 min, followed by 35 cycles of 95 °C for 20 s, 57 °C for 20 s, and 68 °C for 20 s, and a final extension at 68 °C for 60 s.

## Results

### Bioassay and F1 hybrid screening

The difference in the level of chlorantraniliprole resistance between the susceptible strain and the six field populations ranged from 600-fold (low) to 12,500-fold (high), and that of flubendiamide resistance ranged from 428-fold (low) to 129,186-fold (high) (Table 1). The resistance ratios in all populations differed between the two diamide insecticides; for example, Anseong showed higher resistance to chlorantraniliprole, and Cheongju had higher resistance to flubendiamide. However, all six populations exhibited cross-resistance to both diamide insecticides. Similar results and cross-resistance were reported by Cho et al. (2018) in their bioassay using two representative diamide insecticides cyantraniliprole, cyclaniliprole, and tetraniliprole. Among the six populations, Yeoju had the highest resistance ratio to chlorantraniliprole and Gangneung to flubendiamide. Despite its high resistance to both insecticides, the Yeoju population could not recover during the resistance selection test because of its small size at the time of collection. The Jindo population, which had the largest population size at the time of collection, maintained its size even after resistance selection for four subsequent generations for each insecticide. After the fifth generation was exposed to resistance selection, the resistance ratio of the Jindo population was 29,950-fold for chlorantraniliprole and 135,286-fold for flubendiamide, increasing 4.3-fold for chlorantraniliprole and 8.8-fold for flubendiamide compared with that of the 4th generation. This selection system was designated as Diamide resistant (Di-R) strain and used as a basic strain for the resistance mechanism along with the susceptible strain. In the case of F1 hybrids, 12 out of 15 pairs of R♂ × S♀ and eight pairs of S♂ × R♀ hatched successfully; the total number of larvae was higher in R × S combinations (data not shown).

Although the experiments were not sufficiently repeated due to deficiency in populations, there was no statistically significant difference in the resistance of each R × S and S × R combination. The resistance ratio was 10,000-fold for chlorantraniliprole (20–30 mg L^-1^) and 40,000-fold for flubendiamide (30–50 mg L^-1^). Therefore, we conclude that the inheritance of the two diamide-related genes is partially recessive, as reported for other insects (Richardson et al. 2020).

### Phylogenetic relationships

Partial mt genome (approximately 2.5 kb) sequences of the field populations were compared with those of the two susceptible strains, Di-R derived from Jindo population, and four F1 hybrids. There were 22 single nucleotide polymorphisms (SNPs) among the 2,544 nucleotide sequences. The largest sequence variation was detected in the Anseong population; this sequence contained 15 SNPs compared with the reference susceptible strain. The SNP pattern may be different even though samples were collected from different regions of the city, such as from Gangneung and Jindo. Some level of nucleotide sequence diversity was observed between Di-R and the susceptible strain after four generations of resistance selection, but no SNPs were found in Yeoju, Icheon, and Miryang populations. Seven SNPs were identified in the NAS susceptible strain. This strain was used as another reference strain for LAMP and multiplex PCR.

### I4790M mutation and indel in ryanodine receptors

The partial genome sequence analysis of RyR based on cDNA and gDNA revealed no G4946E mutation in the susceptible strain and Di-R, but confirmed the presence of the I4790M mutation in Di-R (Fig. 2A). The PCR using gDNA as a template and RyD_4790UF and RyR_I4790M_R primers further confirmed the presence of only the I4790M mutation. The PCR product size of the susceptible and resistant strains was different (Fig. 2B) and resistant specific indel was confirmed via sequencing. The resistance-specific band obtained using cDNA was similar in size (about 0.5 kb, Fig. 2C) and identical in sequence composition to that of gDNA. The part of intron that was not spliced remained intact. Therefore, it was confirmed that resistance-specific indel exists and is partially transcribed. The resistance-specific indel was verified in gDNA and in cDNA by PCR using BAW_RyR_F3 and B3 primers, the two primers specific for this indel (Fig. 2C).

**Fig. 2.**
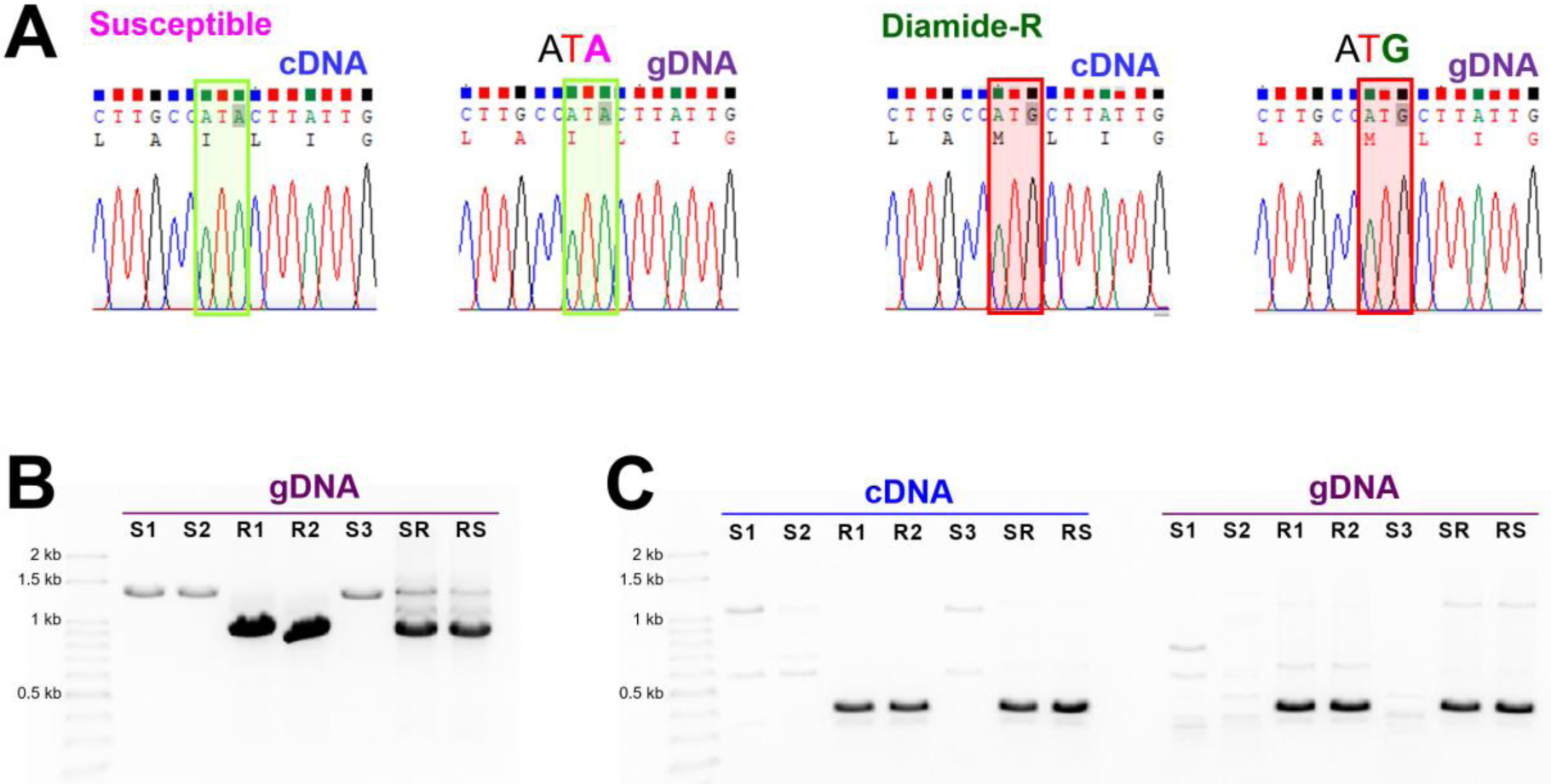
Representative chromatograms of direct sequencing for I4790M mutation genotyping in ryanodine receptors from (A) susceptible and diamide-resistant strains of *Spodoptera exigua*. Green and red boxes denote the I4790M mutation sites in susceptible and resistant strains, respectively. (B) Diamide-resistant strain-specific transcript variants and existence of indels were confirmed via gel electrophoresis of PCR results using gDNA as a template. (C) Resistant alleles were amplified from cDNA and gDNA templates of the diamide resistant strain and its F1 hybrids, SR (male susceptible × female resistant strain) and RS (male resistant × female susceptible strain). F1 hybrids generated by single pair mating using susceptible and diamide-resistant strains. Primer information is documented in Table 2

### LAMP and multiplex PCR

The bioassay identified the I4790M mutation and the resistance-specific indel in all resistant populations (Fig. 3). The intron length of susceptible and resistant strains was 1110 nt and 693 nt, respectively (susceptible, MT292621; Di-R, MT292620), and no SNPs were found between the two susceptible strains. Some SNPs existed among the resistant populations, and a few SNPs were unique to the Anseong population (Fig. 3). The general PCR using F3 and B3 LAMP primers confirmed the presence of resistance-specific traits (Fig. 2C), and then LAMP was performed using four primers (two internal primers, FIP and BIP) (Fig. 4). After 2 h of incubation at 61, 63, and 65 °C, the best primer sensitivity was obtained at 65 and 63 °C, whereas false positives were observed at 61 °C. Therefore, additional LAMP experiments were conducted at 65 °C. To verify the effectiveness of the loop primers LF and LB, the experiments were run using four primers, only LF, only LB, and both LF and LB, and the incubation time was reduced from 2 h to 1 h 30 min (Fig. 5). The amplification efficiency visibly increased with addition of LF, whereas the amplification efficiency of LB was low when compared with that of LF. The addition of a loop primer may reduce the reaction time by 30 min (Fig. 5).

**Table 2.**
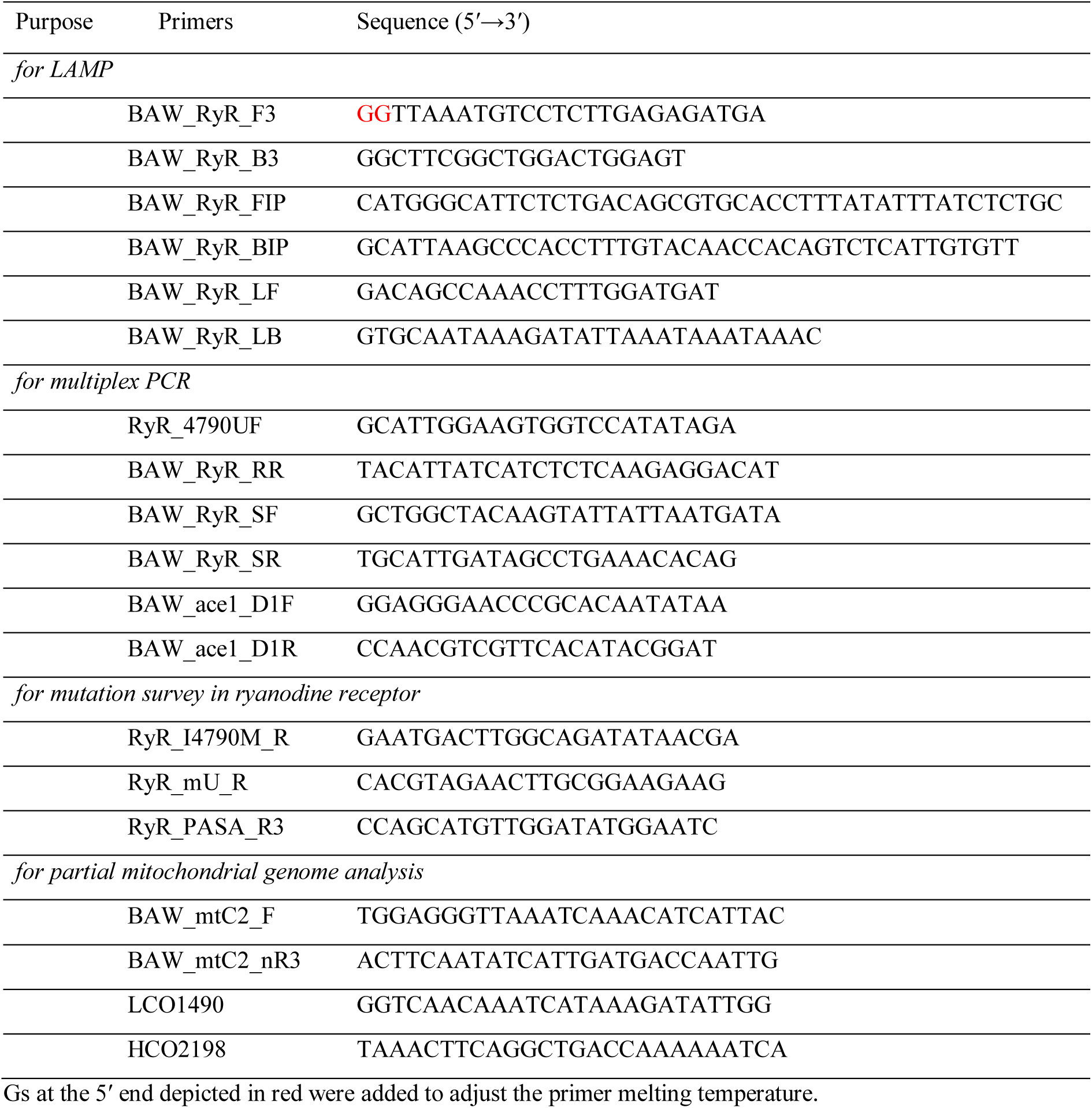
Primers used for lamp loop-mediated isothermal amplification (LAMP), multiplex PCR, and mutation survey in ryanodine receptor

**Fig. 3.**
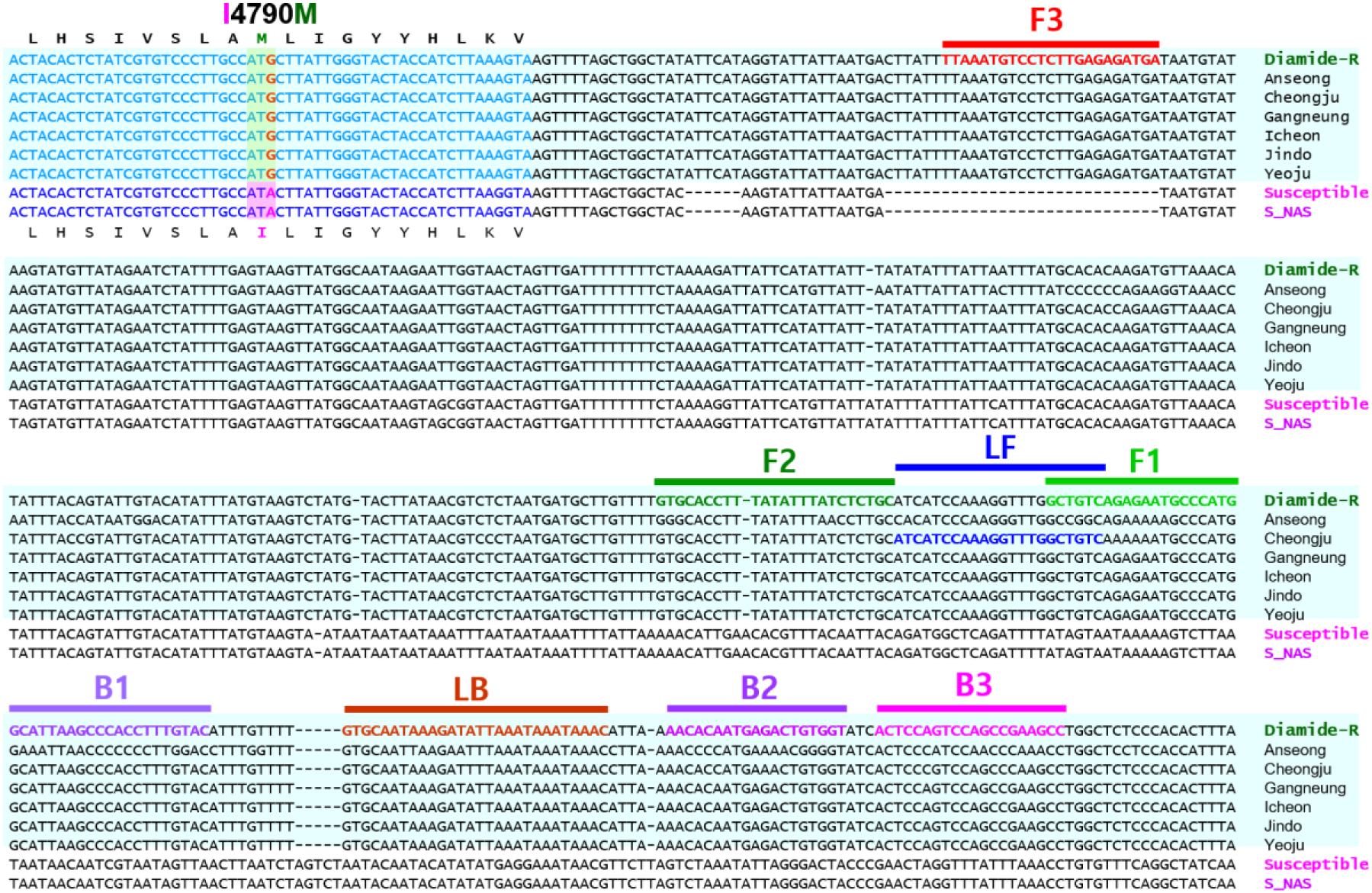
Location of primers and primer binding regions on partial sequences of ryanodine receptors of *Spodoptera exigua.* Sequence alignment of six local populations with two susceptible and a diamide-resistant strain. The I4790M mutation was found in the diamide-resistant strain and six local populations. Amino acid sequences annotated according to the DNA sequence. Lamp loop-mediated isothermal amplification (LAMP) primers designed on the intron region. Internal primer FIP consists of F1c (complementary sequences of F1) and F2. Internal primer BIP is composed of B1 and B2c (complementary sequences of B2). Four essential LAMP primers (F3, FIP, BIP, and B3) generate the dumbbell structure and two loop primers, LF and LB, accelerate the LAMP reaction (see Nagamine et al. 2002 for details). Additional information about primers is documented in Table 2

**Fig. 4.**
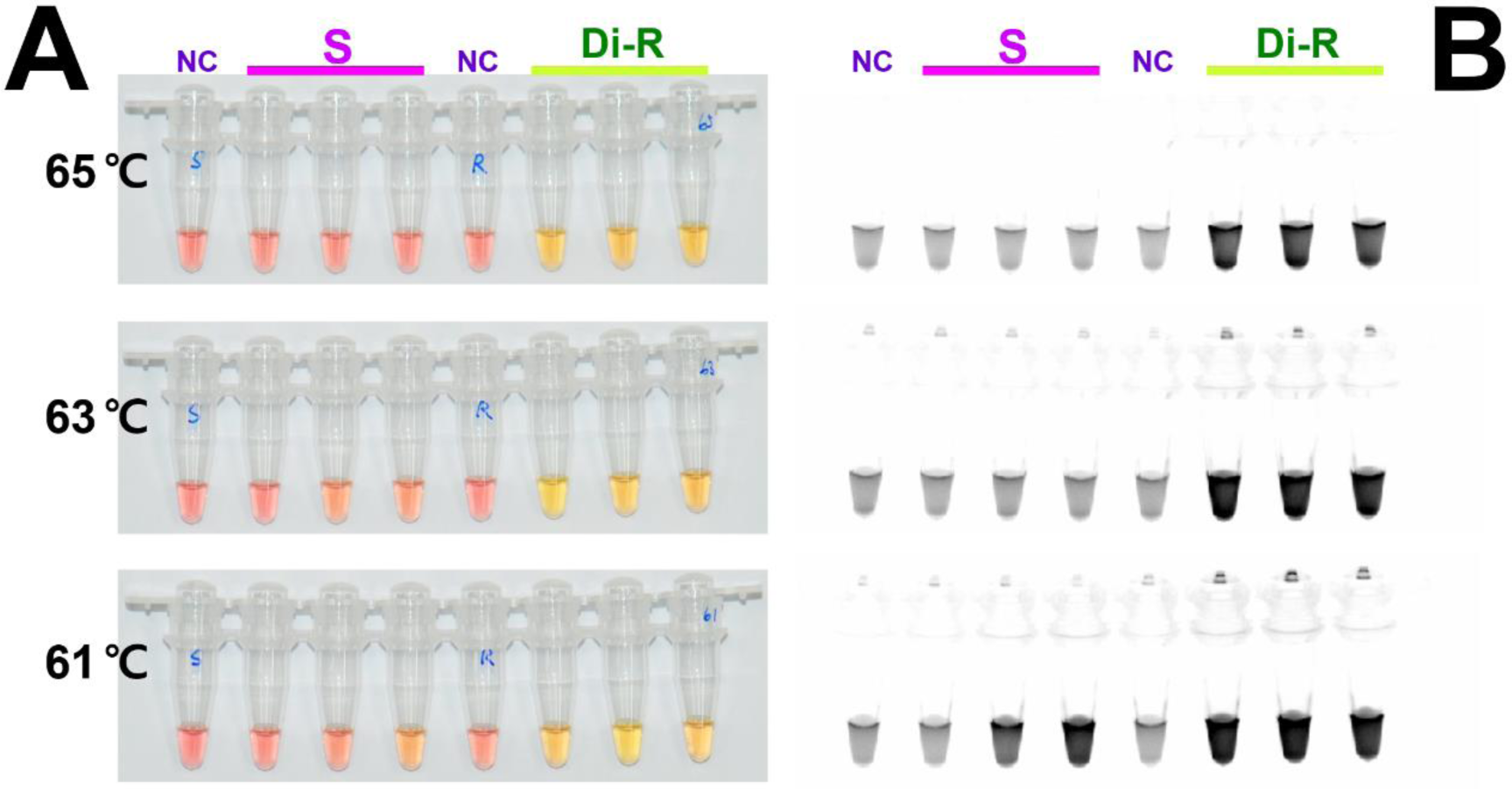
Sensitivity of the lamp loop-mediated isothermal amplification (LAMP) assay results at three temperatures under (A) visible light, (B) ultraviolet light with Cyber Green, and (C) for diamide-resistance allele detection. The sensitivity of the LAMP assay was tested using four primers (BAW_RyR_F3, BAW_RyR_B3, FIP and BIP) at three incubation temperatures, 65, 63 and 61°C for 2 h. The original pink color of the reaction mixture turned yellow in a positive reaction when product was formed but remained pink in negative reactions. NC: negative control; S: susceptible strain; Di-R: diamide-resistant strain

**Fig. 5.**
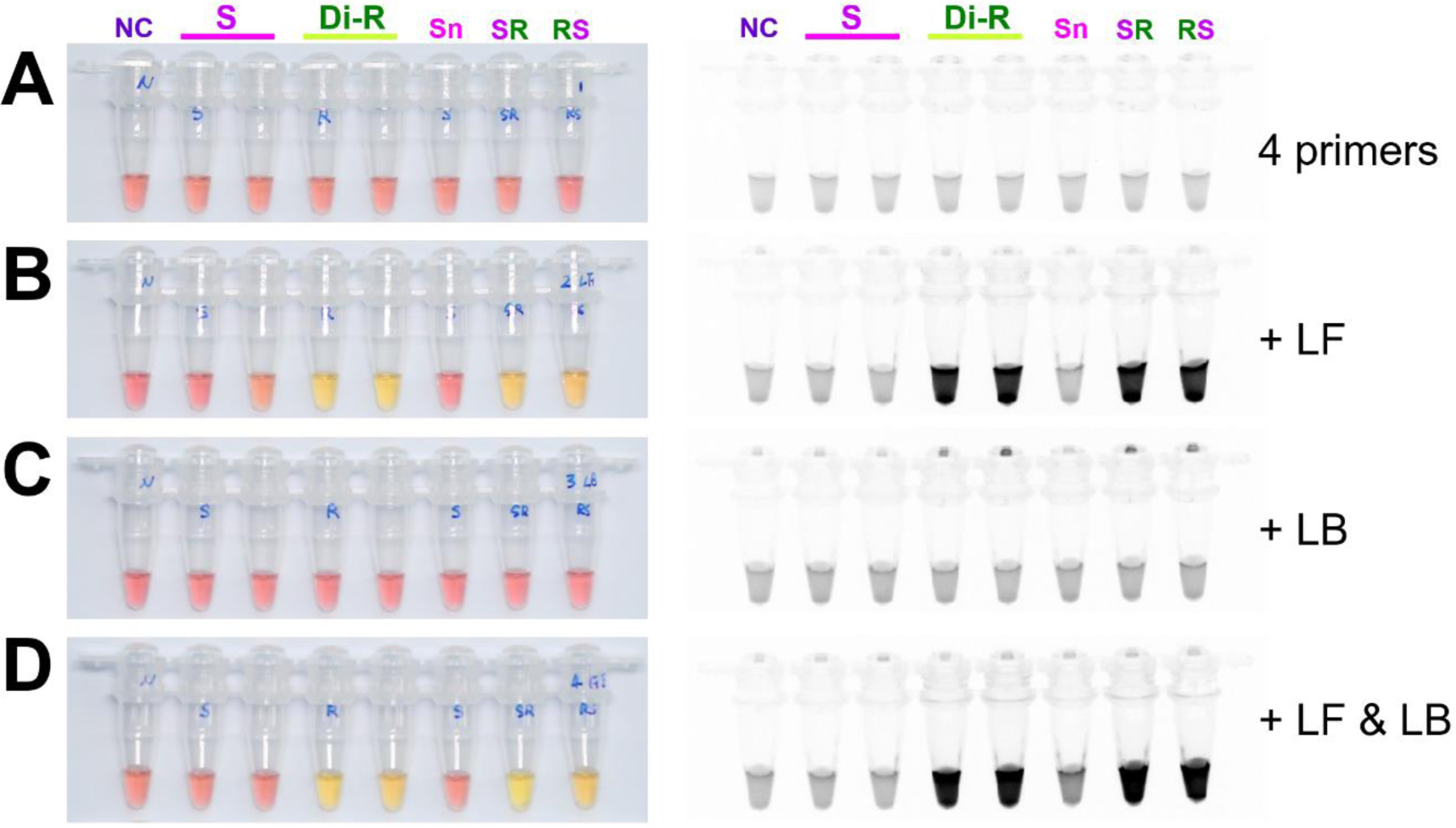
The lamp loop-mediated isothermal amplification (LAMP) assay results with (A) four primers, (B) additional loop primers, (C) loop forward, LF, or (D) loop backward, LB and LF, under visible light (left side) or ultraviolet light with Cyber Green (right side). LAMP assay was tested at incubation temperature of 65°C for 90 min. NC: negative control; S: susceptible strain; Di-R: diamide-resistant strain; Sn: susceptible stain obtained from the National Institute of Agricultural Sciences; SR and RS: F1 hybrid between S and Di-R

To validate the diagnostic concentration limit of LAMP, the initial DNA concentration of 10 ng was decreased 10-fold at each run. Diagnosis was possible for a DNA concentration of at least 100 pg (Fig. 6). Using the DNA releasing technique reported by Kim et al. (2020), sufficient DNA was obtained in the supernatant from larvae and adult tissue after their incubation at 95 ° C for 5 min (Fig. 7). Resistance traits were diagnosed in all 11 populations (Fig. 8). Because LAMP can only confirm the presence of a resistance trait, we developed a multiplex PCR method to detect the heterozygotes. The primer set was selected based on the diagnostic purpose: to diagnose resistance or susceptible traits only, for positive control, or for positive control with either resistance or susceptible traits (Fig. 9).

**Fig. 6.**
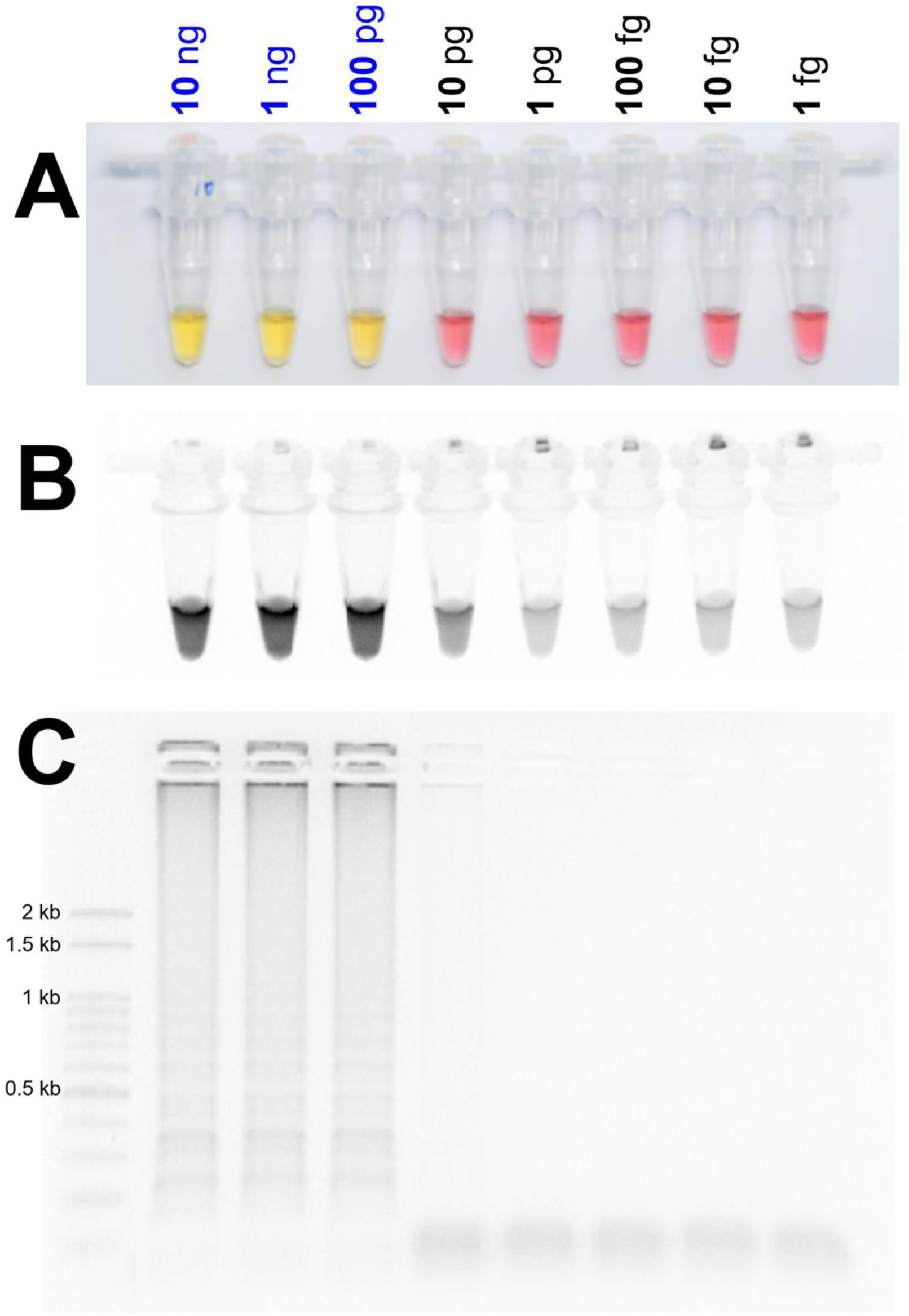
Identification of the detection limit of genomic DNA in the lamp loop-mediated isothermal amplification (LAMP) assay from 10 ng to 1 fg under (A) visible light, (B) ultraviolet light with Cyber Green, and (C) gel electrophoresis. The LAMP assay was tested at an incubation temperature of 65 °C for 90 min with six primers

**Fig. 7.**
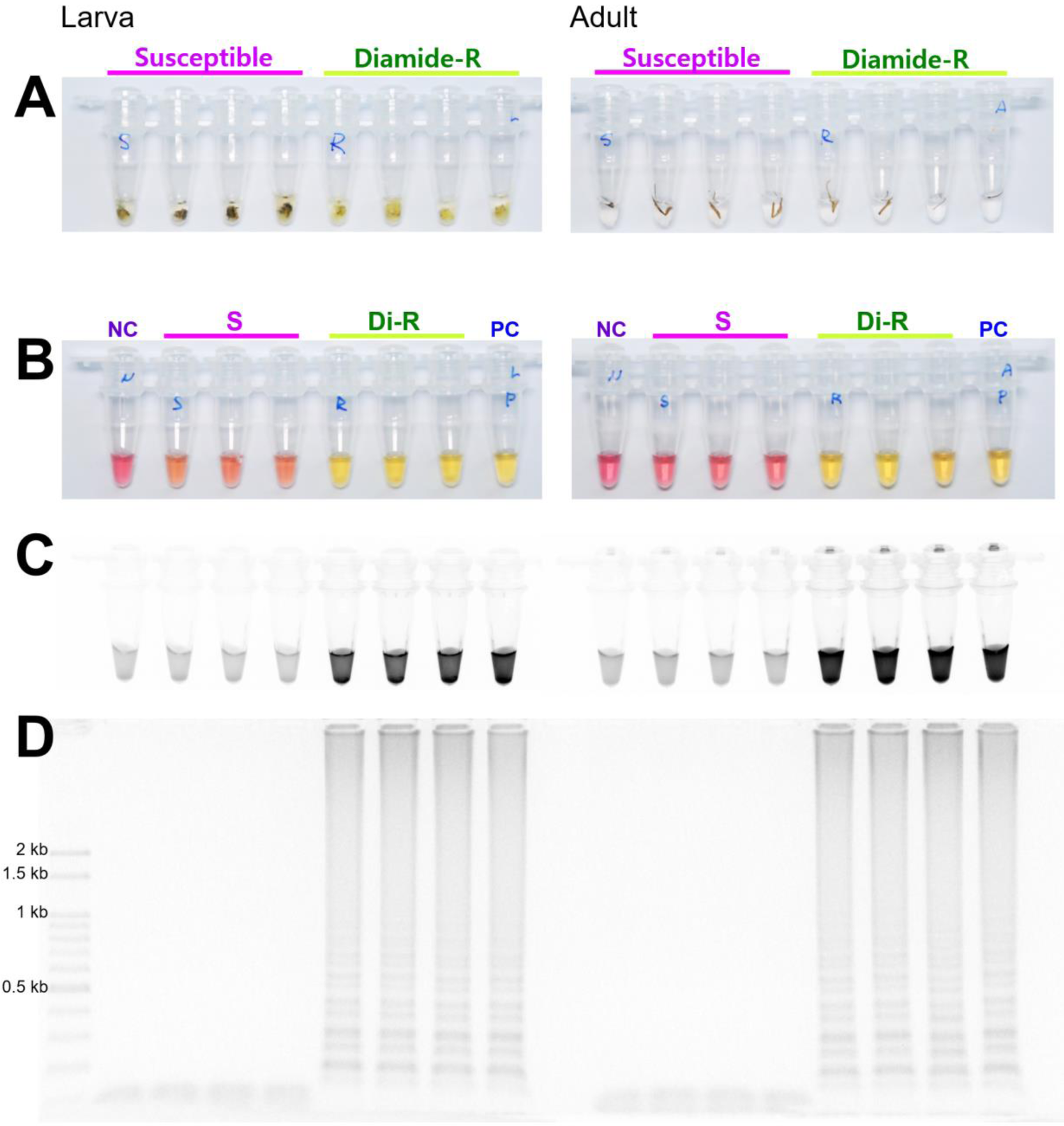
Sensitivity of the lamp loop-mediated isothermal amplification (LAMP) assay results using DNA samples obtained by the DNA releasing technique from insect tissue. (A) About 10 mg of larval tissue or adult leg (or antenna) were incubated in 95 °C for 5 min. LAMP products under (B) visible light, (C) ultraviolet light with Cyber Green, and (d) gel electrophoresis. Abbreviations are as in Fig. 4; PC positive control (isolated DNA from diamide-resistant strain, Di-R)

**Fig. 8.**
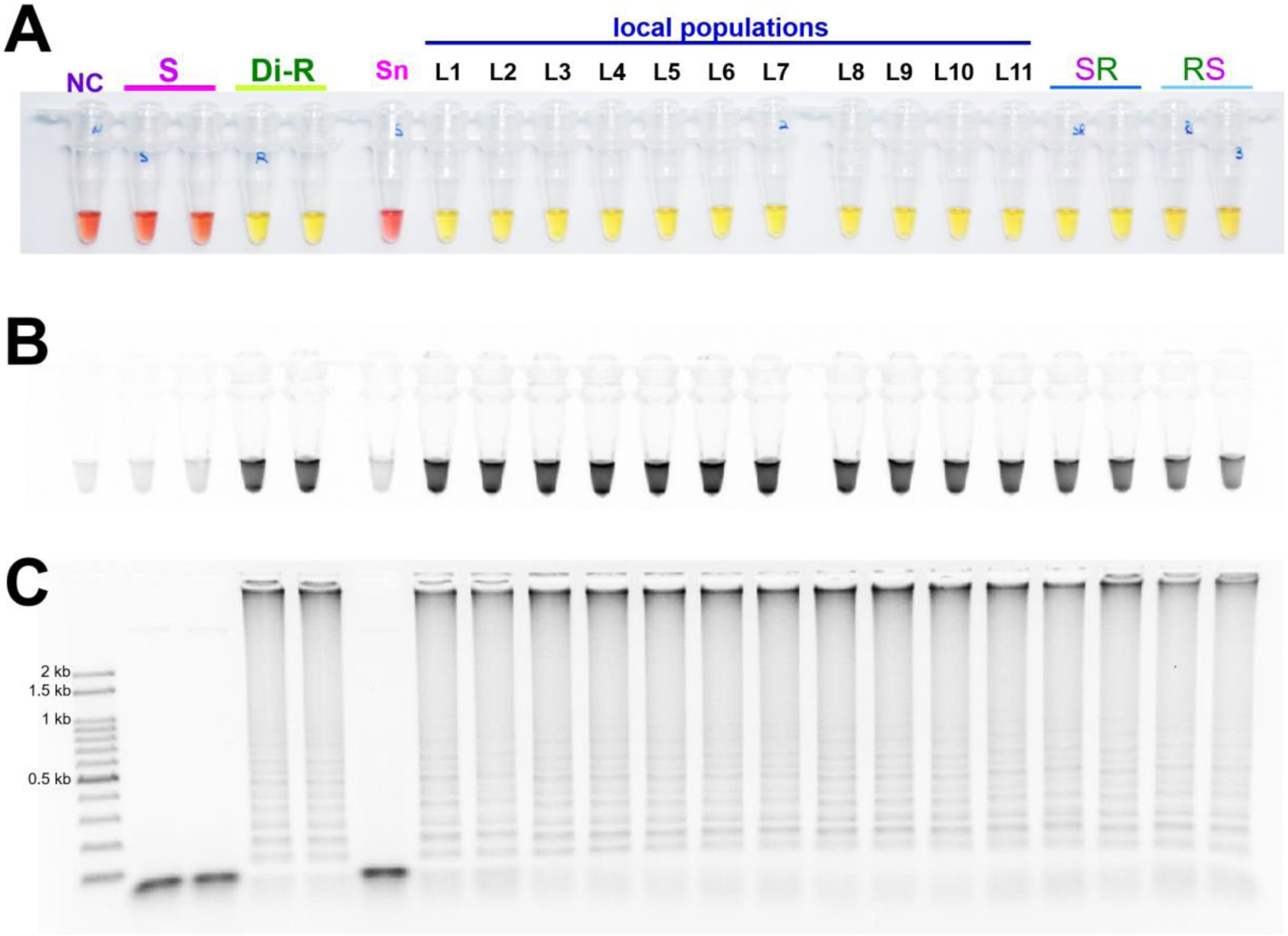
Sensitivity of the lamp loop-mediated isothermal amplification (LAMP) assay results for the resistance allele survey in field-collected local populations with susceptible and diamide-resistant strains under (A) visible light, (B) ultraviolet light with Cyber Green, and (C) gel electrophoresis. NC: negative control; S: susceptible strain; Di-R: diamide resistant strain; Sn: susceptible stain from the National Institute of Agricultural Sciences; SR and RS: F1 hybrids between S and Di-R. L1–L11: field-collected local populations, Anseong (AS), Cheongju (CJ), Gangneung (GN), Icheon (IC), Jindo (JD), Yeoju (YJ), Haenam (HN), Hoengseong (HS), Hongchen (HC), Miryang (MY), and Pyeongchang (PC)

**Fig. 9.**
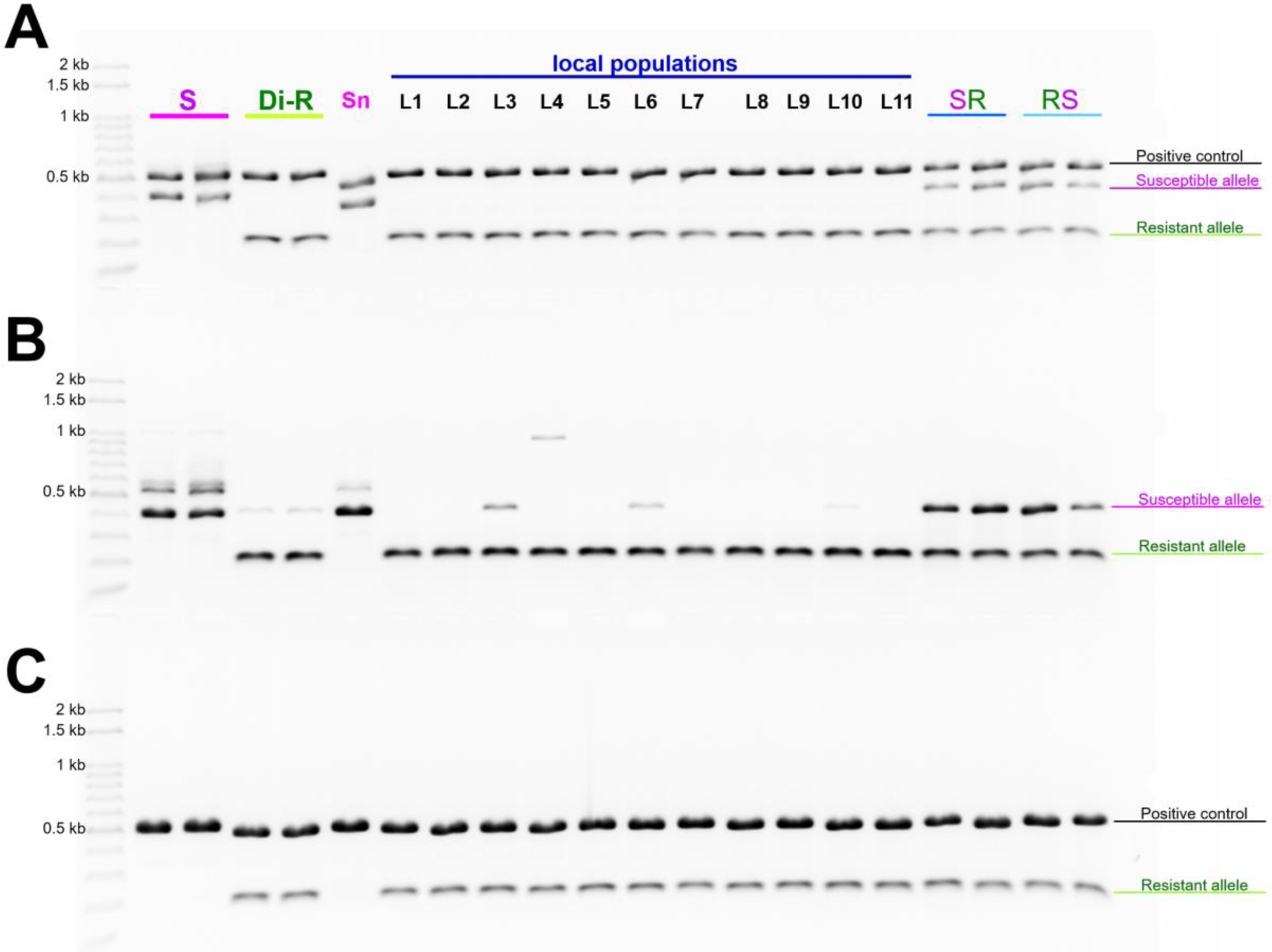
Multiplex PCR results for (A) susceptible and resistance alleles diagnostic combination with positive control or (B) without positive control. *Ace1* type acetylcholinesterase gene was used as positive control. Panel (C) shows the resistant allele possibly amplified with positive control. Abbreviations are as in Fig. 8

## Discussion

Insecticide resistance of *S. exigua*, especially diamide-based, has been reported in many regions worldwide. In Korea, the resistance to diamide in 2014, especially against chlorantraniliprole, was minimal, but 3 years later, highly resistant strains to chlorantraniliprole, over 2,500-fold, were detected in some regions, including Jindo (Cho et al. 2018). We compared the resistance levels reported in 2017 with those in major regions affected by *S. exigua* in 2019 and detected a difference in resistance levels among the regions; the resistance level remained high. Six populations exhibited high cross-resistance to diamide in regions where the management of diamide-based insecticide was exigent. The G4964E mutation in RyR, which is a known diamide resistance mechanism of *S. exigua*, was not identified either in the 2017 populations or in the 2019 populations. This phenomenon has already been reported in *Plutella xylostella*, in which mutations other than G4964 were reported (Guo et al. 2014a). The I4790M mutation was recently identified as a new, major resistance-related mutation in a resistant population of *S. exigua* in China (Zuo et al. 2019) and a population of *S. frugiperda* in Brazil (Boaventura et al. 2020). This mutation was functionally identified via a *Drosophila melanogaster* expression system (Douris et al. 2017). The I4970M mutation was confirmed in Di-R and six populations (Fig. 3), and it was characterized by the presence of a resistance-specific indel in the intron section (Fig. 2). The presence of this indel in the six populations despite their different genetic background, as well as its expression in the intron, indicates that it is closely related to resistance (Fig. 2). However, how this mutation confers resistance is yet to be revealed. G4964E, I4790M, and other mutations in RyR are the main elements of diamide-resistance mechanism in Lepidoptera, with some of them causing over-expression of detoxifying enzymes, such as cytochrome P450s. The transcriptome analysis of resistance mechanisms is currently in progress and is expected to provide insight into the role of the indel in resistance. RyR upregulation has been associated with diamide resistance in *S. exigua* (Sun et al. 2015). However, we detected no difference in RyR expression between resistant and susceptible strains (data not shown). The presence of I4790M mutation and resistance-specific indel was confirmed through expression at cDNA and gDNA levels (Fig. 2C).

Four primers, two internal primers (FIP and BIP) and two resistance-specific indel primers (BAW_RyR_F3, BAW_RyR_B3), used in LAMP successfully diagnosed the resistance-specific marker at 65 °C and 63 °C. Although, LAMP is less sensitive to reaction temperature compared with general PCR, it is important to optimize the reaction conditions for a given primer or other reaction parameters. The highest sensitivity of the LAMP reaction was achieved at 65°C, and addition of a loop primer known to increase the amplification efficiency further improved the diagnostic properties of this protocol. The addition of the loop primer LF reduced the reaction time by about 30 min, suggesting high amplification efficiency of this primer, whereas LB showed a relatively low efficiency (Fig. 5). LAMP was successfully applied to an extensive range of DNA concentrations, from the initial amount of 100 ng to the minimum required DNA concentration of 100 pg when using five primers, four basic primers and LF (Fig. 6). Thus, this protocol is suitable for samples obtained by simple incubation without DNA extraction, in which DNA concentrations are not constant. Sufficient DNA was released by incubation of larval and adult tissues, rendering this method, which requires only a heat block for temperature control, suitable in the field. LAMP, as a time- and labor-efficient protocol, has been utilized in various fields, including ecology, medicine, diagnostic of plant viruses inside insect bodies, and identification of pest species such as *Spodoptera frugiperda* (Lee et al. 2017; Kim et al. 2020), as well as in insecticide resistance allele diagnosis (Badolo et al. 2012, 2015; Choi et al. 2018). Besides LAMP, resistance diagnostics and detection of homotraits or heterotraits can be performed by multiplex PCR and other techniques, but unlike LAMP, these methods can be performed only in a laboratory.

The development of diamide resistance in *S. exigua* is gradually expanding, and resistant strains have been identified in many crops and regions in Korea. According to the Insecticide Resistance Action Committee, insecticide selection is one of the most important factors in resistance management (IRAC 2019). Prior information on the level of resistance distribution is essential for selection of insecticides, especially for diamide resistance, because diamide insecticides are one of the most widely used pesticides in Lepidopteran pest control. In this study, we developed an effective LAMP protocol that can be applied directly in the field without a technical DNA extraction process. Multiplex PCR can be used if a more detailed diagnosis is necessary. A next step is to identify how the resistance-specific indel operates in diamide-resistant *S. exigua*. The presence of indels affects P450 gene expression, such as that of *CYP6g1* in fruit fly (Daborn et al. 2002; Le Goff and Hilliou 2017), or genomic changes associated with insecticide resistance (Faucon et al. 2015; Berger et al. 2016; Duneau et al. 2018; Lucas et al. 2019). Mechanisms responsible for the difference between the RyR mutation and the expression of detoxifying enzyme have been partially revealed (Wang et al. 2016, 2018), but as a kind of long non-coding RNA (lncRNA), there are no reports on the indel in intron and lncRNA expression from *S. exigua*. Functional analysis of this lncRNA will help to elucidate more specific mechanisms of diamide resistance.

## Acknowledgments

This study was supported by the Cooperative Research Program for Agriculture Science & Technology Development (Project No. PJ01358802), the Rural Development Administration, Republic of Korea.

## Declarations

### Funding

This study was funded by Rural Development Administration, Republic of Korea (project number: PJ01358802, project title: Development of molecular diagnostic technique for insecticide resistance against Noctuidae pests).

### Conflicts of interest

The authors declare that they have no conflict of interest.

### Ethics approval

This article does not contain any studies with human participants or animals performed by any of the authors.

### Consent to participate

Not applicable

### Consent for publication

Not applicable

### Availability of data and material

#### Code availability

Not applicable

### Authors’ contributions

J.K., M.K., and G-H.K conceived the study. J.K, J.H.C., and S.R.C. mainly prepared the samples and performed the bioassay. J.K., and H.Y.N. accomplished the LAMP as well as the multiplex PCR method. J.K. and H.Y.N mainly wrote the paper. All authors read and approved the manuscript.

## Notes

### Competing Interest Statement

The authors have declared no competing interest.

## References

Badolo A, Bando H, Traoré A, Ko-Ketsu M, Guelbeogo WM, Kanuka H, Ranson H, Sagnon N, Fukumoto S (2015) Detection of G119S ace-1 (R) mutation in field-collected *Anopheles gambiae* mosquitoes using allele-specific loop-mediated isothermal amplification (AS-LAMP) method. Malar J 14:477. https://doi.org/10.1186/s12936-015-0968-9.

Badolo A, Okado K, Guelbeogo WM, Aonuma H, Bando H, Fukumoto S, Sagnon N, Kanuka H (2012) Development of an allele-specific, loop-mediated, isothermal amplification method (AS-LAMP) to detect the L1014F *kdr-w* mutation in *Anopheles gambiae s.l*. Malar J 11:227. https://doi.org/10.1186/1475-2875-11-227.

Berger M, Puinean AM, Randall E, Zimmer CT, Silva WM, Bielza P, Field LM, Hughes D, Mellor I, Hassani-Pak K, Siqueira HA, Williamson MS, Bass C (2016) Insecticide resistance mediated by an exon skipping event. Mol Ecol 25(22):5692–5704. https://doi.org/10.1111/mec.13882

Boaventura D, Bolzan A, Padovez FEO, Okuma DM, Omoto C, Nauen R (2020) Detection of a ryanodine receptor target-site mutation in diamide insecticide resistant fall armyworm, *Spodoptera frugiperda*. Pest Manag Sci 76:47–54. https://doi.org/10.1002/ps.5505

CABI (2020) Invasive species compendium. https://www.cabiorg/isc/search/index. Accessed 04 May 2020

Cho S-R, Kyung Y, Shin S, Kang W-J, Jung DH, Lee S-J, Park G-H, Kim SI, Cho SW, Kim HK, Koo H-N, Kim GH (2018) Susceptibility of field populations of *Plutella xylostella* and *Spodoptera exigua* to four diamide insecticides. Korean J Appl Entomol 57(1):43–50. https://doi.org/10.5656/KSAE.2018.02.0.009

Cordova D, Benner EA, Sacher MD, Rauh JJ, Sopa JS, Lahm GP, Selby TP, Stevenson TM, Flexner L, Gutteridge S, Rhoades DF, Wu L, Smith RM, Tao Y (2006). Anthranilic diamides: a new class of insecticides with a novel mode of action, ryanodine receptor activation. Pestic Biochem Physiol 84(3):196–214. https://doi.org/10.1016/j.pestbp.2005.07.005

Coronado R, Morrissette J, Sukhareva M, Vaughan DM (1994). Structure and function of ryanodine receptors. Am J Physiol, 266(6):C1485–C1504. https://doi.org/10.1152/ajpcell.1994.266.6.C1485

Daborn PJ, Yen JL, Bogwitz MR, Le Goff G, Feil E, Jeffers S, Tijet N, Perry T, Heckel D, Batterham P, Feyereisen R, Wilson TG, ffrench-Constant RH (2002) A single p450 allele associated with insecticide resistance in *Drosophila*. Science 27;297(5590):2253–2256. https://doi.org/10.1126/science.1074170

Douris V, Papapostolou K-M, Ilias A, Roditakis E, Kounadi S, Riga M, Nauen R, Vontas J (2017) Investigation of the contribution of RyR target-site mutations in diamide resistance by CRISPR/Cas9 genome modification in *Drosophila*. Insect Biochem Mol Biol 87:127–135. https://doi.org/10.1016/j.ibmb.2017.06.013

Duneau D, Sun H, Revah J, San Miguel K, Kunerth HD, Caldas IV, Messer PW, Scott JG, Buchon N (2018) Signatures of insecticide selection in the genome of *Drosophila melanogaster*. G3 (Bethesda). 8(11):3469–3480. https://doi.org/10.1534/g3.118.200537.

Faucon F, Dusfour I, Gaude T, Navratil V, Boyer F, Chandre F, Sirisopa P, Thanispong K, Juntarajumnong W, Poupardin R, Chareonviriyaphap T, Girod R, Corbel V, Reynaud S, David JP (2015) Identifying genomic changes associated with insecticide resistance in the dengue mosquito *Aedes aegypti* by deep targeted sequencing. Genome Res 25(9):1347–1359. https://doi.org/10.1101/gr.189225.115

Foster SP, Denholm I, Rison JL, Portillo HE, Margaritopoulis J, Slater R (2012) Susceptibility of standard clones and European field populations of the green peach aphid, *Myzus persicae*, and the cotton aphid, *Aphis gossypii* (Hemiptera: Aphididae), to the novel anthranilic diamide insecticide cyantraniliprole. Pest Manag Sci 68(4):629–633. https://doi.org/10.1002/ps.2306

Guo L, Liang P, Zhou X, Gao X (2014a) Novel mutations and mutation combinations of ryanodine receptor in a Chlorantraniliprole resistant population of *Plutella xylostella* (L.). Sci Rep 4:6924. https://doi.org/10.1038/srep06924

Guo L, Wang Y, Zhou X, Li Z, Liu S, Pei L, Gao X (2014b) Functional analysis of a point mutation in the ryanodine receptor of *Plutella xylostella* (L.) associated with resistance to chlorantraniliprole. Pest Manag Sci 70(7):1083–1089. https://doi.org/10.1002/ps.3651

Insecticide Resistance Action Committee (2019) IRAC Online. https://www.irac-onlineorg/about/resistance/mechanisms/. Accessed 01 May 2020

Jeanguenat A (2013) The story of a new insecticidal chemistry class: the diamides. Pest Manag Sci 69(1):7–14. https://doi.org/10.1002/ps.3406

Kim J, Nam HY, Kwon M, Kim H, Yi HJ, Hänniger S, Unbehend S, Heckel, DG (2020) Development of a simple and accurate molecular tool for *Spodoptera frugiperda* species identification using LAMP. bioRxiv https://doi.org/10.1101/2020.04.07.029678

Kumar S, Stecher G, Tamura K (2016) MEGA7: Molecular Evolutionary Genetics Analysis Version 7.0 for bigger datasets. Mol Biol Evol 33(7):1870–1874. https://doi:10.1093/molbev/msw054

Lahm GP, Cordova D, Barry JD (2009) New and selective ryanodine receptor activators for insect control. Bioorg Med Chem 17(12):4127–4133. https://doi.org/10.1016/j.bmc.2009.01.018

Lai T, Li J, Su J (2011) Monitoring of beet armyworm *Spodoptera exigua* (Lepidoptera: Noctuidae) resistance to Chlorantraniliprole in China. Pestic Biochem Physiol 101:198–205. https://doi.org/10.1016/j.pestbp.2011.09.006

Lee PL (2017). DNA amplification in the field: move over PCR, here comes LAMP. Mol Ecol Resour 17(2):138–141. https://doi:10.1111/1755-0998.12548

Le Goff G, Hilliou F (2017) Resistance evolution in *Drosophila* : the case of CYP6G1. Pest Manag Sci 73(3):493–499. https://doi.org/10.1002/ps.4470

Liu JB, Li FY, Dong JY, Li YX, Zhang XL, Wang YH, Xiong LX, Li ZM (2018) Anthranilic diamides derivatives as potential ryanodine receptor modulators: synthesis, biological evaluation and structure activity relationship. Bioorg Med Chem 26(12):3541–3550. https://doi.org/10.1016/j.bmc.2018.05.028

Lucas ER, Miles A, Harding NJ, Clarkson CS, Lawniczak MKN, Kwiatkowski DP, Weetman D, Donnelly MJ, Anopheles gambiae 1000 Genomes Consortium (2019) Whole-genome sequencing reveals high complexity of copy number variation at insecticide resistance loci in malaria mosquitoes. Genome Res 29(8):1250–1261. https://doi.org/10.1101/gr.245795.118

Nauen R (2006) Insecticide mode of action: return of the ryanodine receptor. Pest Manag Sci 62(8):690–692. https://doi.org/10.1002/ps.1254

Nauen R, Steinbach D (2016) Resistance to diamides in epidopteran pests. In: Horowitz A, Ishaaya I (eds) Advances in insect control and resistance management. Springer, Dordrecht, pp 219–240.

Richardson EB, Troczka BJ, Gutbrod O, Davies TGE, Nauen R (2020) Diamide resistance: 10 years of lessons from lepidopteran pests. J Pest Sci. 93:911–928. https://doi.org/10.1007/s10340-020-01220-y

Roditakis E, Vasakis E, Grispou M, Stavrakaki M, Nauen R, Gravouil M, Bassi A (2015) First report of *Tuta absoluta* resistance to diamide insecticides. J Pest Sci 88(1):9–16. https://doi.org/10.1007/s10340-015-0643-5

Sanderson MJ, Wojciechowski MF (2000) Improved bootstrap confidence limits in large-scale phylogenies, with an example from Neo-Astragalus (Leguminosae). Syst Biol 49(4):671–685. https://doi:10.1080/106351500750049761

Sattelle DB, Cordova D, Cheek TR (2008) Insect ryanodine receptors: molecular targets for novel pest control chemicals. Invert Neurosci 8(3):107–119. https://doi.org/10.1007/s10158-008-0076-4

Sparks TC, Nauen R (2015) IRAC: mode of action classification and insecticide resistance management. Pestic Biochem Physiol 121:122–128. https://doi.org/10.1016/j.pestbp.2014.11.014

Steinbach D, Gutbrod O, Luemmen P, Matthiesen S, Schorn C, Nauen R (2015) Geographic spread, genetics and functional characteristics of ryanodine receptor based target-site resistance to diamide insecticides in diamondback moth, *Plutella xylostella*. Insect Biochem Mol Biol 63:14–22. https://doi.org/10.1016/j.ibmb.2015.05.001

Sun L, Qiu G, Cui L, Ma C, Yuan H (2015) Molecular characterization of a ryanodine receptor gene from *Spodoptera exigua* and its upregulation by Chlorantraniliprole. Pestic Biochem Phys 123:56–63. https://doi.org/10.1016/j.cbd.2018.07.001

Sun Z, Xu H (2019) Ryanodine receptors for drugs and insecticides: an overview. Mini Rev Med Chem 19(1):22–33. https://doi.org/10.2174/1389557518666180330112908

Tohnishi, M, Nakao H, Furuya T, Seo A, Kodama H, Tsubata K, Fujioka S, Kodama H, Hirooka T, Nishimatsu T (2005) Flubendiamide, a novel insecticide highly active against lepidopterous insect pests. J Pestic Sci 30(4):354–360. https://doi.org/10.1584/jpestics.30.354

Troczka B, Zimmer CT, Elias J, Schorn C, Bass C, Davies TGE, Field LM, Williamson MS, Slater R, Nauen R (2012) Resistance to diamide insecticides in diamondback moth, *Plutella xylostella* (Lepidoptera: Plutellidae) is associated with a mutation in the membrane-spanning domain of the ryanodine receptor. Insect Biochem Mol Biol 42(11):873–880. https://doi.org/10.1016/j.ibmb.2012.09.001

Troczka BJ, Williams AJ, Williamson MS, Field LM, Lüemmen P, Davies TE (2015) Stable expression and functional characterization of the diamondback moth ryanodine receptor G4946E variant conferring resistance to diamide insecticides. Sci Rep 5:14680. https://doi.org/10.1038/srep14680

Wang X, Chen Y, Gong C, Yao X, Jiang C, Yang Q (2018) Molecular identification of four novel cytochrome P450 genes related to the development of resistance of *Spodoptera exigua* (Lepidoptera: Noctuidae) to chlorantraniliprole. Pest Manag Sci 74(8):1938–1952. https://doi.org/10.1002/ps.4898

Wang XG, Gao XW, Liang P, Shi XY, Song DL (2016) Induction of Cytochrome P450 Activity by the Interaction of Chlorantraniliprole and Sinigrin in the *Spodoptera exigua* (Lepidoptera: Noctuidae). Environ Entomol 45(2):500–507. https://doi.org/10.1093/ee/nvw007

Zuo YY, Wang H, Xu Y, Huang J, Wu S, Wu Y, Yang Y (2017) CRISPR/Cas9 mediated G4946E substitution in the ryanodine receptor of *Spodoptera exigua* confers high levels of resistance to diamide insecticides. Insect Biochem Mol Biol 89:79–85. https://doi.org/10.1016/j.ibmb.2017.09.005

Zuo YY, Ma HH, Lu WJ, Wang XL, Wu SW, Nauen R, Wu YD, Yang YH (2019) Identification of the ryanodine receptor mutation I4743M and its contribution to diamide insecticide resistance in *Spodoptera exigua* (Lepidoptera: Noctuidae). Insect Sci. https://doi.org/10.1111/1744-7917.12695

